# Unified sampling framework and experimental benchmarking of sequence- and structure-based protein models

**DOI:** 10.64898/2026.05.08.723784

**Authors:** Aviv Spinner, Pascal Notin, Samuel Berry, Dana Cortade, Zach Sisson, Svetlana Ikonomova, David Ross, Debora Marks

## Abstract

Generative models are increasingly used for protein design, but the lack of standardized evaluation frameworks limits comparison across model classes and hinders translation to experimental success. Here, we introduce a unified sampling and benchmarking framework that enables controlled sequence generation across alignment, protein language, and structure-based models, and apply it to Tobacco etch virus (TEV) protease. Across hundreds of thousands of designed sequences, different models explore distinct regions of sequence space with no clear computational selection metrics to assess enzymatic function. Experimental evaluation reveals large differences in functional outcomes, ranging from non-functional variants to sequences with 9-fold higher activity than wildtype. Machine learning-designed libraries achieve a 39.32% hit rate (percentage of variants matching or exceeding wildtype activity) compared to 6.06% for an error-prone PCR baseline. Structure-based models perform best overall, with hit rates of 74.4% and 66.8% for ESM-IF1 and ProteinMPNN, respectively. Commonly used selection metrics do not strongly correlate with experimental activity, highlighting a gap between in silico evaluation and enzyme function. Together, these results establish a generalizable framework for benchmarking generative protein models and demonstrate the necessity of experimental validation for guiding model development and sequence prioritization.

## 1. Introduction

Generative artificial intelligence (AI) is rapidly transforming biology, enabling new approaches to protein design by leveraging both natural sequence data and structural information. Familiar examples of successful AI such as GPT (Brown et al., 2020) demonstrate that models trained on large-scale datasets can generalize to a wide range of downstream tasks and perform high quality generation. Motivated by this success, computational biologists have increasingly integrated machine learning into studies of protein function and design (Notin* et al., 2024).

Natural selection provides billions of years of evolutionary experiments, encoding fundamental principles of protein structure and function within sequence space. Protein engineering seeks to harness these principles, but remains time-intensive and costly when performed experimentally at scale (Dane Wittrup, 2012; Alberghina, 2000). As a result, machine learning approaches have been introduced to predict the effects of mutations and suggest new variants with improved or altered function (Madani & al., 2023; Shin et al., 2021; Lian et al., 2022; Schiff et al., 2024; Sumida et al., 2024; Thadani et al., 2023). In particular, the field of protein variant effect prediction has produced large benchmarking datasets and general evaluation guidelines, demonstrating that different models perform variably across tasks (Notin et al., 2023; Livesey et al., 2024). Recent community efforts, such as protein engineering challenges and tournaments (Armer et al., 2024; van Niekerk et al., 2026), further highlight both the promise of these approaches and the need for standardized evaluation practices.

Despite these advances, the field lacks methodological consensus for using machine learning to generate protein sequences. Existing studies are highly bespoke, differing in training data, model architectures, sampling strategies, filtering criteria, and success metrics. As a result, comparisons between generative models remain difficult, and many designed proteins are non-functional, leading to wasted experimental effort. While prior work has explored limited comparisons between generative models (Anonymous, 2024; Darmawan et al., 2023; Johnson et al., 2024; Pillutla et al., 2022; Spinner et al., 2024), there is currently no standard-ized framework for evaluating and visualizing differences across arbitrary model classes.

In this work, we introduce a unified sampling framework and a rigorous benchmarking strategy for protein design models. Our framework enables consistent sequence generation across diverse model classes, including alignment-based, structure-based, and protein language models. We apply this framework to Tobacco etch virus (TEV) protease, a well-characterized and widely used enzyme, as a proof-of-concept system. TEV protease is a 236 amino acid cysteine protease that canonically cleaves the sequence ENLYFQ|S and serves as a robust model for studying substrate-protease interactions.

Using a diverse suite of generative models, we generated hundreds of thousands of TEV variants under controlled sampling conditions. We then performed large-scale computational and experimental benchmarking to evaluate generated sequence diversity, model-specific biases, and enzymatic function. This work establishes a generalizable framework for comparing generative protein models and provides a foundation for integrating experimental validation to develop predictive filtering metrics for protein design (Figure 1).

**Figure 1.**
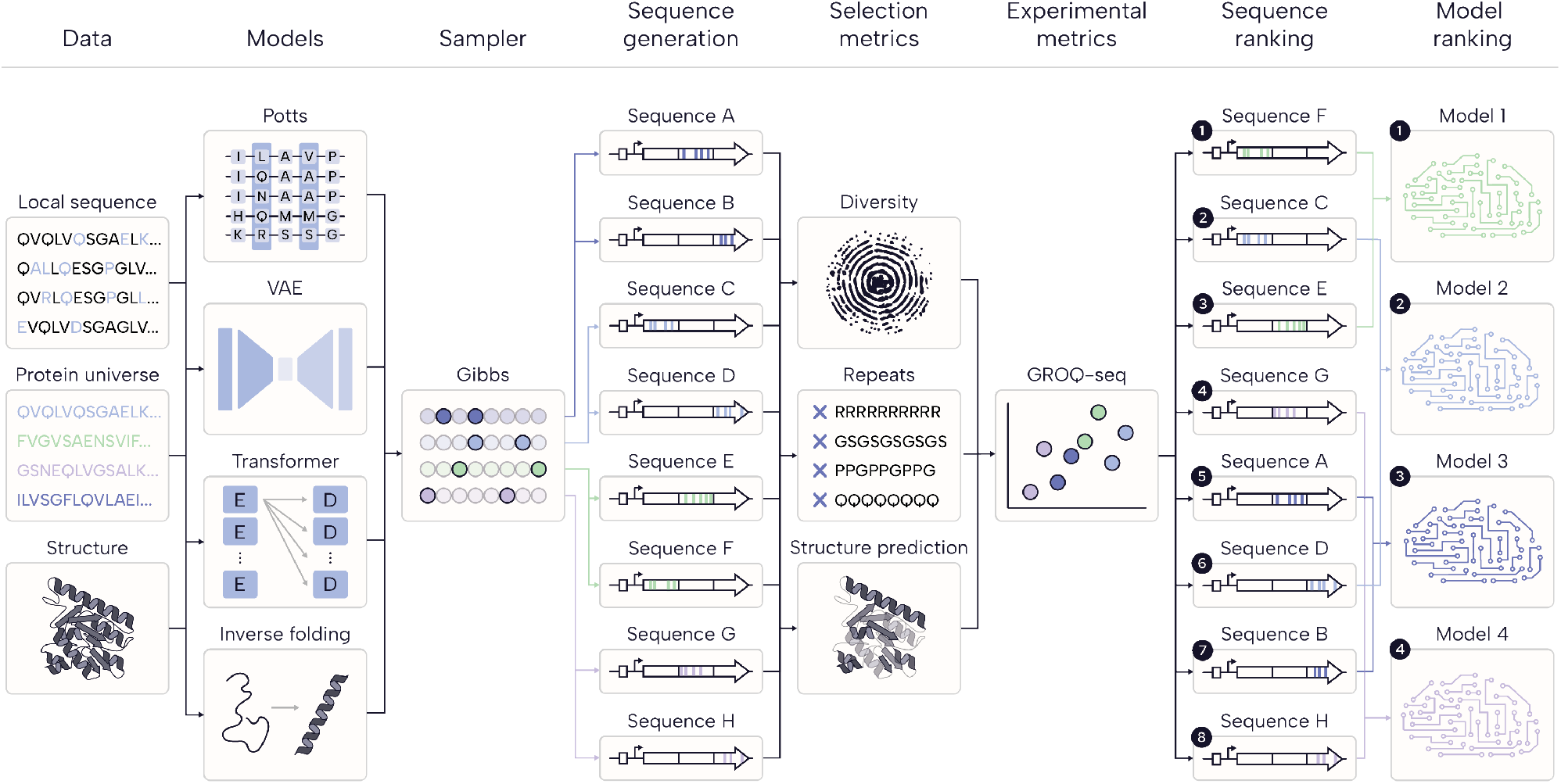
Schematic of the benchmarking pipeline integrating diverse training data (local sequence alignments, protein universe, and structure) and model classes (Potts/PSSM, VAE, transformer-based language models, and inverse folding models). Sequences are generated using a consistent Gibbs sampling procedure, enabling controlled comparison across models. Generated variants are evaluated using selection metrics and experimentally assayed using GROQ-seq. Sequence- and model-level rankings are then derived, providing a unified framework for comparing generative strategies in protein design.

We summarize our key contributions as follows:

- A unified sampling framework for controlled sequence generation across alignment-based, autoregressive, and structure-conditioned protein models;
- A large-scale experimental benchmark of ∼13k designed TEV variants with paired model provenance and quantitative functional measurements, released as a community resource;
- A systematic empirical analysis of commonly used selection metrics, showing that while individual metrics capture modest signal, no single score reliably prioritizes functional variants, and identifying filtering as a critical and under-explored bottleneck in ML-guided protein design;
- A direct cross-model comparison under controlled sampling conditions, demonstrating that structure-conditioned models (ProteinMPNN (Dauparas et al., 2022), ESM-IF1 (Hsu et al., 2022)) consistently out-perform sequence- and language-based models in functional enrichment.

## 2. Methods

### 2.1. Overview of Models and Sampling Framework

We selected a diverse set of generative models spanning multiple data modalities and architectural classes (Figure 1), including alignment-based models (Position specific scoring matrix [PSSM, implemented as in (Hopf & Marks, 2017)] and Potts/EVCouplings (Marks et al., 2011)), variational autoencoders (EVE (Frazer et al., 2021)), protein language models (ESM2 (Lin et al., 2023), Tranception), and structure-based inverse folding models (ESM-IF1 (Hsu et al., 2022), ProteinMPNN (Dauparas et al., 2022)). Each model presents unique generation strategies; however, to enable direct comparison, we created a unified Python framework that applies a consistent Gibbs sampling procedure across all models.

This framework allows any model capable of computing pseudo log-likelihoods over sequences to be used for controlled sequence generation. All code is available at: https://github.com/pascalnotin/protein_sampling.

### 2.2. Training Data

Two primary sources of unsupervised training data were used: multiple sequence alignments (MSAs) and protein structure.

To construct the MSA, we performed five iterations of JackHMMER (Johnson et al., 2010) against UniRef100 (Suzek et al., 2007), MGnify (Mitchell et al., 2020), and BFD (Tunyasuvunakool et al., 2021) with a relative bitscore threshold of 0.2. Given the high substrate specificity of TEV protease, we prioritized a smaller, functionally relevant alignment over a larger but more diverse one. Full-length sequences were retrieved using Easel (S.R. Eddy, unpublished) and realigned with ClustalOmega (Sievers et al., 2011). Columns not corresponding to the query sequence were removed, resulting in an alignment of 4,748 sequences with no more than 20% gaps per position. This alignment defines the set of homologous sequences used for downstream in silico scoring and comparison.

For structure-based models, we generated a full-length structure of TEV protease using AlphaFold3 (Abramson et al., 2024; Tunyasuvunakool et al., 2021). Because experimentally resolved structures often exclude the C-terminal region due to autocatalysis (Abramson et al., 2024; Kapust et al., 2001), we modeled the protease in complex with its substrate to obtain a complete and functionally relevant structure.

### 2.3. Model Training

Alignment-based models (PSSM, EVCouplings, EVE) were trained with a sequence reweighting threshold of *θ* = 0.9 to downweight highly similar sequences. Structure-based models (ESM-IF1, ProteinMPNN) were used with default settings. Protein language models (ESM2, Tranception) were used without fine-tuning.

### 2.4. Sequence Generation via Gibbs Sampling

All models generated sequences using Gibbs sampling to ensure methodological consistency. Gibbs sampling iteratively mutates one position at a time by sampling amino acids according to model-derived likelihoods, allowing controlled exploration of sequence space while preserving sequence length.

We implemented a Gibbs sampling method for each model, extending the approach introduced by Berry et al. (Berry et al., 2026), which focused on alignment-based models (e.g., Potts/PSSM), to additional model classes used here including variational autoencoders, autoregressive protein language models, and structure-based inverse folding models. This extension required adapting the Gibbs sampling procedure to accommodate differences in model likelihood formulations and conditioning mechanisms across architectures.

We included a linear distance restraint to control the number of mutations in sampled sequences. This restraint linearly penalizes mutations away from the reference sequence and rewards mutations that restore it, with its magnitude defined in the same units as the model likelihood. Because likelihood scales differ across models, restraint values are not directly comparable and were instead calibrated empirically to achieve comparable mutational distances from wildtype.

Each sampling step consists of 236 position-wise updates corresponding to the length of TEV protease. For each model, we performed 25 sampling steps across 32 parallel chains, discarding the first step of each chain. This yielded 768 sequences per sampling condition.

Two key hyperparameters controlled generation: a distance penalty and temperature. We selected model-specific distance restraints to yield approximately five mutations from wildtype on average, enabling balanced comparison across models (Table A2).

### 2.5. Experimental Constraints on Sequence Design

To ensure compatibility with downstream DNA synthesis, the TEV protease sequence was divided into three segments of 78 amino acids. Mutations were restricted within each segment, while all other positions were fixed to wildtype by assigning probability 1. The S219V mutation was fixed across all sequences to prevent autocatalysis.

### 2.6. Filtering

Minimal filtering was applied to preserve sequence diversity and minimize bias introduced at this step. Sequences were retained if they contained a complete catalytic triad (H46/D81/C151) and had ≤ 10 mutations from wildtype.

### 2.7. Codon Optimization

Gene variants were designed by introducing specified amino acid substitutions into a codon-optimized wildtype sequence defined in the original GROQ-seq TEV protease assay (Spinner et al., 2026).

### 2.8 Assembly and Cloning

Variants were assembled with a barcoded plasmid backbone using a 5-part Golden Gate assembly (Cortade et al., 2026). The circuit utilizes inducible promoters from the Marionette system (Meyer et al., 2019).

### 2.9 GROQ-seq TEV Protease Assay

Quantitative fitness was measured using the GROQ-seq TEV protease assay (Spinner et al., 2026). The assay was developed as part of the GROQ-seq platform (Cortade et al., 2024).

## 3. Results

### 3.1. Generation and Filtering

Across all models, parameter combinations, and three sections of the protein, this procedure generated 338,688 total sequences. After filtering, 71,032 sequences remained, of which 67,664 were unique (Table 1). These sequences were used for all downstream analyses.

**Table 1.** Sequence counts across generation, filtering, and testing. All models generated 48,384 sequences prior to filtering. Counts are not deduplicated across columns and may overlap between models (Dedup: deduplicated sequences; Active: intact catalytic triad; ≤10: ≤ 10 mutations from wildtype; Test: number of variants experimentally assayed). ProteinMPNN produces the largest number of unique sequences, while PSSM and Potts are the only models for which all sequences contain a complete active site.

| MODEL | DEDUP. | ACTIVE | $\leq 10$ | TEST |
| --- | --- | --- | --- | --- |
| PSSM | 9990 | 9990 | 9648 | 2209 |
| POTTS | 9923 | 9923 | 9608 | 2213 |
| EVE | 13476 | 13352 | 11798 | 3012 |
| ESM2 | 9916 | 9915 | 8747 | 1789 |
| TRANCEPTION | 10861 | 9012 | 8406 | 2445 |
| ESM-IF1 | 11329 | 10067 | 9843 | 2049 |

### 3.2. Computational Comparisons Across Models

We quantified sequence diversity using several complementary metrics:

- Total positions mutated across all generated sequences
- Hamming distance from wildtype
- Distance to the nearest sequence in the alignment
- Pairwise distances within generated sequences
- Mutation frequency within the substrate-binding region (Kapust et al., 2001)

Distinct patterns emerged across model classes (Table A1). Alignment-based models (PSSM, Potts) produced highly similar mutational landscapes, while models such as EVE and Tranception explored a broader set of positions. Protein language models, particularly ESM2, tended to generate sequences with a higher number of mutations from wildtype.

Interestingly, the average distance from wildtype closely matched the distance to the nearest alignment sequence, suggesting that generated sequences remain within the natural sequence manifold despite controlled diversification.

Differences were also observed in how models mutate the substrate-binding region. The substrate-binding region is defined based on prior structural work (Kapust et al., 2001) and includes residues {46, 51, 67, 146, 148, 169, 170, 171, 174, 176, 178, 209, 211, 213, 214, 216, 218, 220}. Some models (e.g., ESM2, ProteinMPNN) introduced mutations in this region more frequently, while others (e.g., EVE) largely avoided it, reflecting differing inductive biases learned during training.

To visualize mutational preferences of generation, we constructed a multidimensional scaling (MDS) plot (Armer et al., 2024) of all generated sequences that were experimentally tested (Figure 2A). These analyses revealed qualitatively distinct textures of sequence space. Each model occupies a distinct region, with clustering patterns reflecting underlying training data and inductive biases. Models trained on similar data (e.g., alignment-based models) cluster closely, while those trained on different modalities (e.g., structure-vs. language-based models) explore divergent regions. Notably, EVE exhibited the widest spread in sequence space, indicating more aggressive exploration.

**Figure 2.**
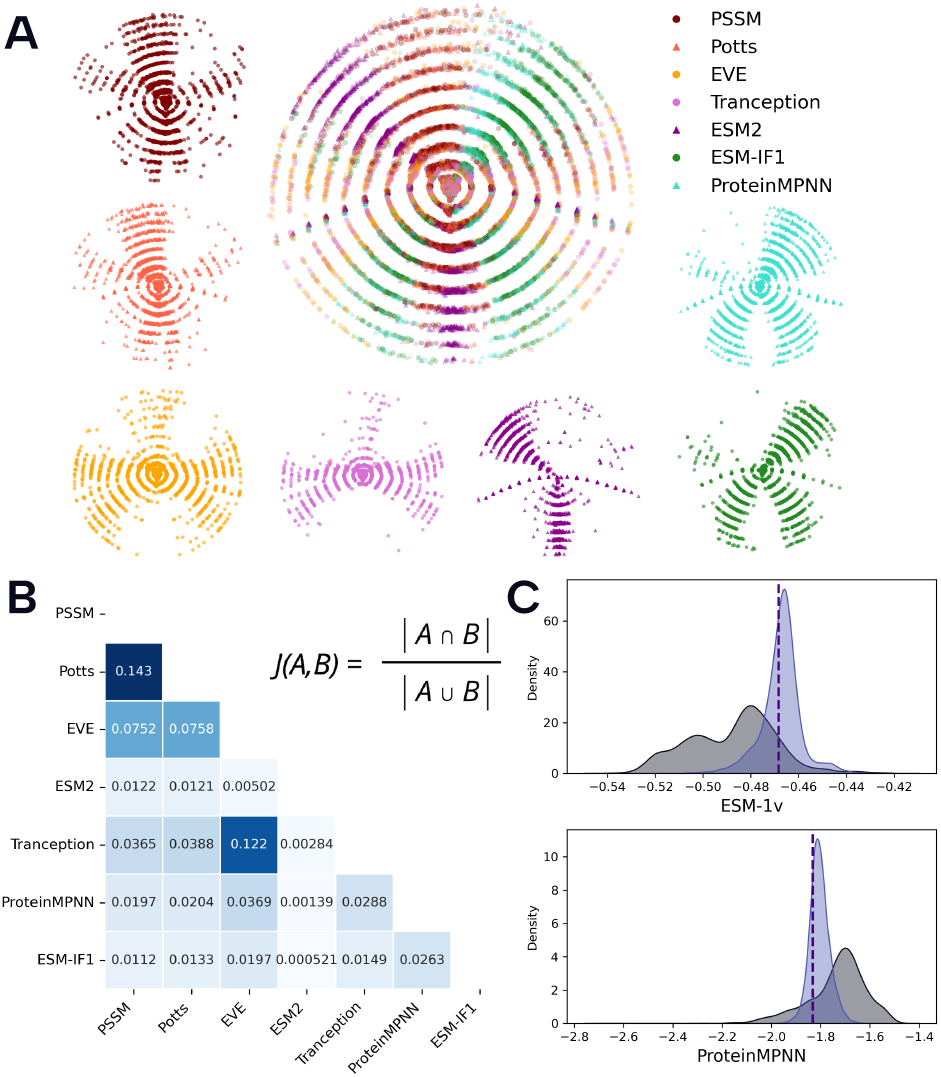
Sequence space structure and computational evaluation of generated variants across models. (A) Multidimensional scaling (MDS) of generated sequences based on pairwise Hamming distance. (B) Pairwise Jaccard similarity between sets of generated sequences. (C) Distributions of fitness prediction scores (pseudo log-likelihoods from ESM-1v and ProteinMPNN, as in (Johnson et al., 2024)) for generated sequences (purple) compared to homologous sequences (gray). Model-specific breakdowns are shown in Supplementary Figure S1.

Pairwise comparisons using the Jaccard similarity index further highlighted relationships between models (Figure 2B). PSSM and Potts exhibited near-identical generation behavior, while overlap between EVE and Tranception suggested shared characteristics despite differences in architecture.

Finally, we evaluated generated sequences using two principal selection metrics from the COMPSS framework (Johnson et al., 2024), ESM-1v and ProteinMPNN, and contextualized these scores relative to the TEV multiple sequence alignment (Figure 2C). For both metrics, sequences from all models clustered within a relatively narrow and favorable range compared to the broader distribution of homologous sequences from the training MSA, likely reflecting the mutational distance constraints imposed during generation. This enrichment suggests that our unified sampling framework preferentially produces sequences that satisfy established computational filters.

Among the models tested, structure-based methods performed strongest by these criteria: ESM-IF1 yielded the highest fraction of sequences passing the ESM-1v threshold, while ProteinMPNN-derived sequences achieved the most favorable ProteinMPNN scores among sequences that passed this filter (Supplementary Figure S1).

We additionally implemented the computational filtering strategy proposed in COMPSS as a reference point (Supplementary Figure S2). Summary statistics across all selection metrics introduced in (Johnson et al., 2024) and (Spinner et al., 2024) are reported in Table A3.

### 3.3. Experimental Measurements Differ from Computational Predictions

To directly assess how well selection metrics reflect functional outcomes, we compared a broad panel of sequence- and structure-based scores to experimentally measured pro-tease activity (Figure 3). We analyzed two experimental conditions: S12, which provides the largest overall dynamic range for assessing functional variation, and S20, which offers improved resolution among top-performing variants.

**Figure 3.**
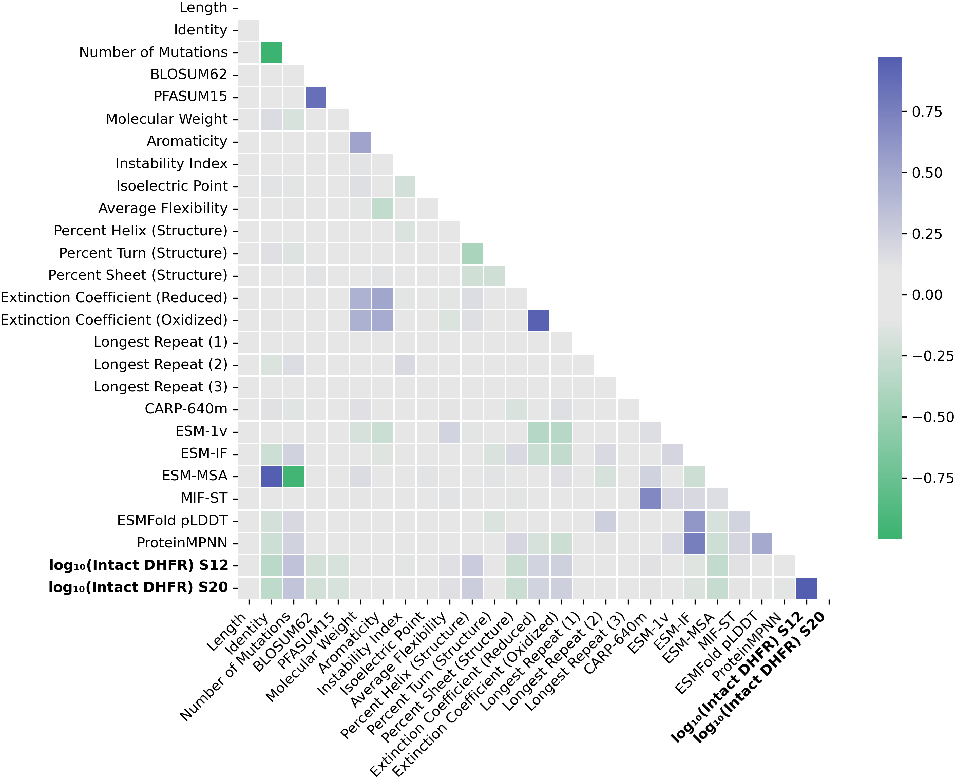
Limited resolution of common selection metrics for filtering functional variants. Modest pairwise Spearman correlations are observed between sequence- and structure-based features, model-derived scores, and experimentally measured protease activity. Correlation patterns are consistent across S12 and S20, which are themselves strongly correlated, but differ in resolution: S12 captures broad activity variation, while S20 better resolves top-performing variants. Across both conditions, most selection metrics show limited association with function, highlighting the limitations of heuristic filtering for prioritizing candidates.

Across all metrics, correlations with experimental activity were generally modest, indicating that while these metrics capture some signal, they have limited standalone predictive power for prioritizing variants. Notably, because lower assay values correspond to higher protease activity, the observed positive correlation with mutation count (i.e., more mutations correlating with lower measured values, and thus higher activity) is expected. The strongest associations with experimental data were observed for the number of mutations (Spearman *ρ* = 0.326) and predicted helical content (Spearman *ρ* = 0.264), as well as a negative correlation with the ESM-MSA prediction (Spearman *ρ* = *™*0.28), though all relationships remained modest.

These results underscore that, despite their convenience, commonly used selection metrics provide limited resolution for prioritizing variants within large candidate pools, and high-quality experimental measurements remain the definitive benchmark for evaluating protein function. Given that these metrics are typically used as a filtering step following sequence generation, this highlights a critical and under-explored bottleneck in ML-guided protein design.

### 3.4. Experimental Comparisons Across Models

We experimentally evaluated a subset of designed TEV protease variants and observed a wide range of functional outcomes, spanning from completely non-functional sequences to variants exhibiting activity up to ∼9-fold higher than the reference wildtype. All model classes produced high-performing sequences, including variants exceeding wild-type activity (Figure 4A; Supplementary Figure S4). At the level of individual variants, substantial differences were observed in the best-performing sequences across models (Supplementary Figure S4).

**Figure 4.**
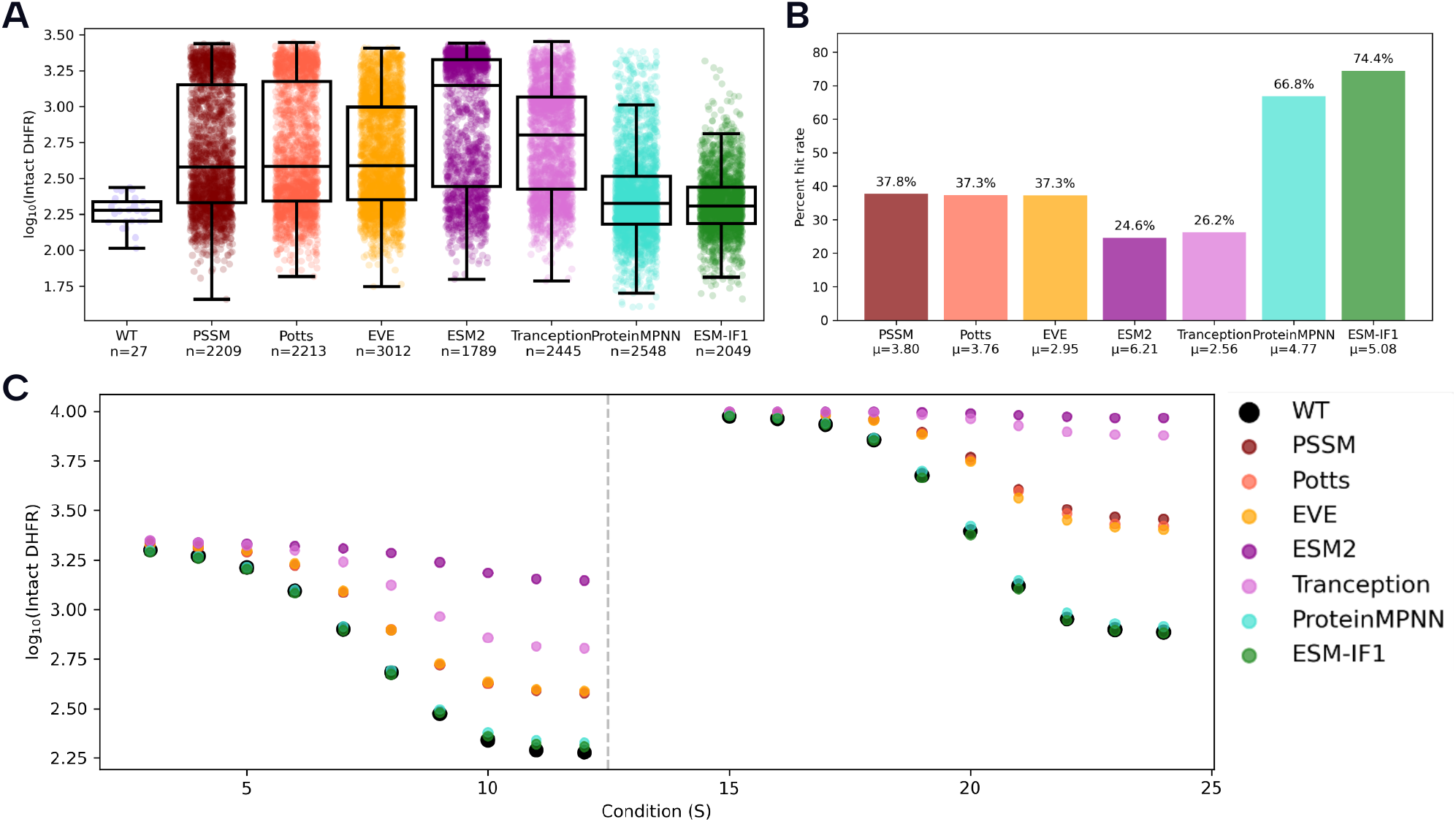
Experimental evaluation of designed variants across models. (A) Distribution of activity (log_10_(Intact DHFR), lower = higher protease activity) for all tested variants, pooled across conditions and stratified by model (*n* = sequences per model). All models generate functional variants spanning a wide activity range. (B) Hit rate (fraction of sequences at or above the highest wildtype replicate; *µ* = average mutations per sequence). Structure-based models (ESM-IF1, ProteinMPNN) achieve the highest hit rates, compared to 6.06% for the epPCR baseline (Supplementary Figure S3). (C) Median activity across 20 experimental conditions. Relative performance is consistent across conditions, with structure-based models outperforming sequence- and language-based approaches.

ProteinMPNN produced the highest-activity variant, achieving an approximately nine-fold improvement over wildtype. The top variants from Tranception and ESM-IF1 exhibited comparable gains of ∼5 to 6-fold, while ESM2, EVE, PSSM, and Potts yielded variants with more modest but still significant improvements of ∼3 to 4-fold. These results highlight that while all model classes are capable of generating functional improvements, structure-based and hybrid approaches are more likely to produce extreme outliers with substantially enhanced activity.

To systematically compare performance across models and conditions, we evaluated all sequences under 20 experimental conditions (Figure 4C) and defined a “hit rate” metric. Specifically, a sequence was considered a hit if its measured activity was at or better than the wildtype sequence with the highest DHFR value, effectively using the worst-performing wildtype replicate as a conservative functional threshold (Figure 4A). This definition enables consistent comparison across experimental conditions.

Using this metric, machine learning-designed libraries substantially outperformed a naive mutagenesis baseline. Compared to an error-prone PCR (epPCR) library, which yielded 6.06% of sequences at or above the wildtype threshold, the ML-designed libraries achieved an average hit rate of 39.32%, despite having comparable mutational loads (Supplementary Figure S3). This control indicates that model-guided sequence generation enriches for functional variants far more effectively than random mutagenesis.

We next examined performance across different model classes. Clear differences emerged between sequence-based, language-based, and structure-based approaches. Structure-based models performed best overall, with hit rates of 74.4% and 66.8% for ESM-IF1 and ProteinMPNN, respectively (Figure 4B). Alignment-based models (PSSM, Potts, and EVE) and protein language models (ESM2 and Tranception) also substantially outperformed the epPCR baseline, achieving hit rates approximately four-to six-fold higher than random mutagenesis, but showed lower overall enrichment than the structure-based models.

Across experimental conditions, model-specific performance trends were largely consistent, though absolute activity levels varied depending on assay conditions (Figure 4C). This indicates that while experimental conditions influence measured activity, the relative ranking of model performance is stable across conditions.

## 4. Discussion

We developed a unified framework for sampling and bench-marking diverse generative protein models and applied it to TEV protease to enable controlled comparisons across model classes. Standardized generation coupled with large-scale experimental validation reveals that models explore distinct regions of sequence space and differ substantially in their ability to produce functional variants. Structure-based approaches consistently yield the highest enrichment for function and the strongest individual variants, although all models generate some improved sequences.

One possible contributing factor is that models such as ProteinMPNN are trained to decode sequences in arbitrary order, which allows efficient conditioning on fixed residues while sampling only a subset of positions. This flexibility aligns naturally with the Gibbs sampling procedure used here, enabling more effective conditional sequence generation under structural constraints. In contrast, selection metrics show limited ability to distinguish model performance or predict experimental outcomes. Together, these findings highlight the importance of model inductive biases in shaping functional sequence generation and underscore the central role of experimental validation in assessing generative protein design.

Selection metrics, as currently used to filter generated sequences, provide limited power for distinguishing model performance or prioritizing variants in practice. Though not sufficient for reliable prioritization on their own, they represent a meaningful but under-leveraged signal within the filtering step. These metrics are typically deployed as heuristic filters applied after sequence generation, yet this filtering stage has received far less systematic study than generative modeling itself. For instance, the distributions of scores from metrics such as ESM-1v and ProteinMPNN are substantially tighter than the spread observed in experimental measurements, making it difficult to resolve meaningful differences between models using *in silico* evaluations alone.

Although (Johnson et al., 2024) represents the closest available baseline for computational triage of designed sequences, it was not designed to formally rank generative models. We therefore use it as an informative point of comparison rather than a definitive evaluation framework. While it generally ranks model classes in a similar order, it does not fully recapitulate the distinctions observed through experimental data. Notably, although structure-based models perform best under these computational filters, this advantage is modest relative to the much larger separation observed experimentally.

This limitation is further underscored by the modest relationships between selection metrics and experimental outcomes. Across a broad set of sequence- and structure-based scores (Figure 3), most metrics show only modest individual correlations with measured protease activity. This lack of concordance highlights the difficulty of relying on any single score as a predictor of functional performance. However, modest global correlations do not preclude strong predictive performance in specific regimes; for example, certain thresholds may reliably identify non-functional variants even if overall rank-ordering remains imperfect.

At the same time, the improved experimental performance of structure-based models suggests that incorporating structural constraints provides a meaningful inductive bias, particularly in this system where substrate-bound information is explicitly included during generation.

Our results, together with recent work (Johnson et al., 2024), suggest that improving the filtering step—through better metrics, calibrated thresholds, or learned combinations of metrics—may be as important as advances in generative modeling for practical protein design. Rather than relying on individual metrics, a promising direction is to systematically combine and calibrate these partially informative signals, potentially via an additional layer of machine learning, to improve prediction, filtering, and ranking of designed sequences. Such approaches will likely require large experimental datasets across multiple proteins to disentangle context-dependent relationships between metrics and diverse protein functions. Importantly, extending these analyses beyond a single enzyme to multiple protein families will be critical for determining whether these relationships generalize and for developing robust, broadly applicable strategies for model evaluation and sequence prioritization.

## 5. Conclusion

We present a unified framework for sampling and bench-marking generative protein models that enables direct comparisons across diverse model classes. Using this framework, we generated over 300,000 TEV variants from seven different models. Model choice strongly shapes both the regions of sequence space explored and the likelihood of pro-ducing functional proteins, with structure-based approaches providing the most consistent enrichment for activity.

More broadly, our results underscore a critical gap between computational evaluation and experimental reality. Existing metrics are insufficient to resolve meaningful differences in model performance, reinforcing the need for experimental benchmarking as a central component of model assessment. The framework introduced here provides a scalable foundation for such efforts and establishes a path toward integrating experimental data to develop more predictive evaluation strategies. As generative models continue to evolve, systematic, experimentally grounded comparisons will be essential for translating advances in AI into reliable tools for protein design.

## Acknowledgements

The Align Foundation is supported by Griffin Catalyst and Schmidt Sciences.

## Disclaimer

Certain commercial equipment, instruments, or materials are identified in this paper in order to specify the experimental procedure adequately. Such identification is not intended to imply recommendation or endorsement by the National Institute of Standards and Technology, nor is it intended to imply that the materials or equipment identified are necessarily the best available for the purpose.

## A. Supplementary Tables

**Table A1.** Summary of mutational diversity and substrate-binding-region perturbation across models. “Mutated positions” is the number of unique residue positions mutated across all generated sequences. “Hamming WT” is the average Hamming distance from wildtype, and “Hamming MSA” is the average distance to the nearest sequence in the alignment. “Min pairwise dist.” is the minimum pairwise distance within generated sequences. The final two columns specifically quantify mutations in the substrate-binding region: the percentage of sequences with at least one substrate-binding-region mutation and the percentage of all mutations that occur in substrate-binding-region residues.

| Model | Mutated positions | Hamming WT | Hamming MSA | Min pairwise dist. | Substrate-binding region Seq. w/ mut. | Mut. fraction |
| --- | --- | --- | --- | --- | --- | --- |
| PSSM | 190 | 4.114 | 4.114 | 1.926 | 38.4% | 14.5% |
| Potts | 187 | 4.130 | 4.130 | 1.949 | 38.5% | 14.4% |
| EVE | 232 | 4.220 | 4.220 | 2.302 | 23.9% | 7.6% |
| ESM2 | 143 | 6.153 | 6.153 | 1.605 | 46.2% | 14.6% |
| Tranception | 232 | 3.634 | 3.634 | 2.468 | 28.6% | 11.4% |
| ESM-IF1 | 139 | 5.095 | 5.095 | 1.472 | 40.8% | 8.8% |
| ProteinMPNN | 189 | 4.996 | 4.996 | 1.747 | 45.8% | 14.2% |

**Table A2.** Hyperparameters used for Gibbs sampling across models. Temperatures denote sampling stochasticity, while distance restraints control deviation from the wildtype sequence during generation.

| Model | Temperatures | Distance restraints |
| --- | --- | --- |
| PSSM | 0.1, 0.2, 0.3, 0.4, 0.5, 0.6, 0.7 | 1.5, 2, 2.5 |
| Potts | 0.1, 0.2, 0.3, 0.4, 0.5, 0.6, 0.7 | 1.5, 2, 2.5 |
| EVE | 0.1, 0.2, 0.3, 0.4, 0.5, 0.6, 0.7 | 2.5, 3, 3.5 |
| ESM2 | 0.1, 0.2, 0.3, 0.4, 0.5, 0.6, 0.7 | 3, 3.5, 4, 4.5 |
| Tranception | 0.1, 0.2, 0.3, 0.4, 0.5, 0.6, 0.7 | 3.5, 4, 4.5 |
| ESM-IF1 | 0.1, 0.2, 0.3, 0.4, 0.5, 0.6, 0.7 | 3, 3.5, 4 |
| ProteinMPNN | 0.1, 0.2, 0.3, 0.4, 0.5, 0.6, 0.7 | 3, 3.5, 4 |

**Table A3.** Average selection metrics for all generated sequences. Values are averaged across all generated sequences for each model.

| Metric | PSSM | Potts | EVE | ESM2 | Tranception | ESM-IF1 | ProteinMPNN |
| --- | --- | --- | --- | --- | --- | --- | --- |
| Length | 236 | 236 | 236 | 236 | 236 | 236 | 236 |
| Identity | 0.9826 | 0.9825 | 0.9821 | 0.9739 | 0.9846 | 0.9784 | 0.9788 |
| BLOSUM62 | -0.7057 | -0.6974 | -0.9722 | -0.8673 | -1.3896 | -0.6290 | -0.8649 |
| PFASUM15 | -0.2347 | -0.2181 | -0.4506 | -0.1812 | -0.8621 | 0.2148 | -0.2112 |
| Molecular weight | 26794.24 | 26794.37 | 26820.35 | 26846.64 | 26766.03 | 26805.55 | 26797.96 |
| Aromaticity | 0.1051 | 0.1051 | 0.1063 | 0.1071 | 0.1038 | 0.1053 | 0.1053 |
| Instability index | 36.2091 | 36.1909 | 36.2822 | 35.9068 | 36.1313 | 36.8575 | 36.6037 |
| Isoelectric point | 8.8108 | 8.8150 | 8.8409 | 8.8466 | 8.8639 | 8.6983 | 8.8355 |
| Average flexibility | 0.9984 | 0.9984 | 0.9973 | 0.9982 | 0.9982 | 0.9986 | 0.9979 |
| Percent helix | 0.1262 | 0.1262 | 0.1235 | 0.1255 | 0.1239 | 0.1145 | 0.1193 |
| Percent turn | 0.2274 | 0.2274 | 0.2290 | 0.2261 | 0.2297 | 0.2304 | 0.2314 |
| Percent sheet | 0.3775 | 0.3776 | 0.3784 | 0.3824 | 0.3756 | 0.3791 | 0.3805 |
| Extinction coef. reduced | 32065.03 | 32081.16 | 32629.59 | 32657.12 | 29856.20 | 29896.46 | 30862.64 |
| Extinction coef. oxidized | 32299.72 | 32315.34 | 32871.61 | 32895.68 | 30083.63 | 30109.12 | 31074.26 |
| Longest repeat (1) | -3.0059 | -3.0061 | -3.0103 | -3.0106 | -3.0115 | -3.0088 | -3.0026 |
| Longest repeat (2) | -1.9585 | -1.9618 | -1.9225 | -1.9205 | -1.9688 | -1.9239 | -1.7239 |
| Longest repeat (3) | -1.0025 | -1.0026 | -1.0053 | -1.0027 | -1.0052 | -1.0079 | -1.0032 |
| CARP-640m | -0.3069 | -0.3070 | -0.3302 | -0.3053 | -0.3355 | -0.3188 | -0.3253 |
| ESM-1v | -0.4677 | -0.4677 | -0.4719 | -0.4723 | -0.4681 | -0.4631 | -0.4645 |
| ESM-IF | -1.5358 | -1.5354 | -1.5618 | -1.5235 | -1.5700 | -1.5079 | -1.5049 |
| ESM-MSA | -0.1607 | -0.1607 | -0.1905 | -0.2269 | -0.1794 | -0.2108 | -0.2101 |
| MIF-ST | -0.2515 | -0.2518 | -0.2755 | -0.2487 | -0.2801 | -0.2633 | -0.2632 |
| ESMFold pLDDT | 74.8413 | 74.8478 | 74.5486 | 75.0451 | 74.2012 | 75.0247 | 75.2165 |
| ProteinMPNN | -1.8147 | -1.8149 | -1.8328 | -1.8030 | -1.8364 | -1.7974 | -1.7858 |

## Supplementary Figures

**Figure S1.**
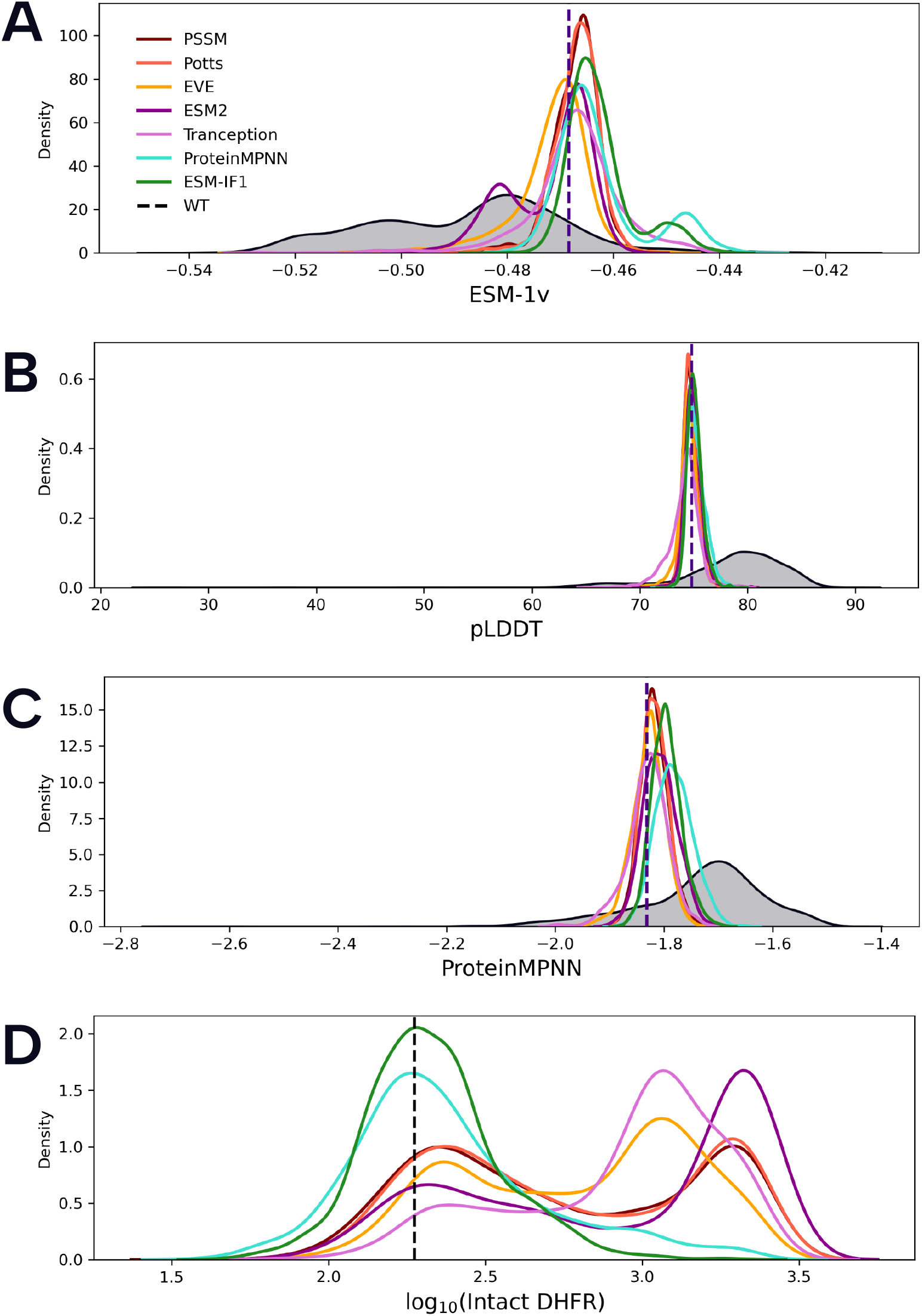
Comparison of selection metrics and experimental outcomes across models. (A–C) Distributions of ESM-1v, ESMFold pLDDT, and ProteinMPNN scores span a narrow range across models, indicating limited separation by selection metrics. Natural homologous sequences (gray) are shifted toward lower scores for sequence-based metrics (ESM-1v) but toward higher scores for structure-based metrics (pLDDT and ProteinMPNN). Across metrics, most designed sequences cluster at or above the wildtype (dashed line), reflecting enrichment under model-guided sampling. ProteinMPNN scores are most favorable for sequences generated by ProteinMPNN itself, although differences between models remain modest. (D) Experimental activity (log_10_(Intact DHFR), where lower values indicate higher protease activity) exhibits a substantially broader distribution, revealing greater functional diversity than suggested by *in silico* scores.

**Figure S2.**
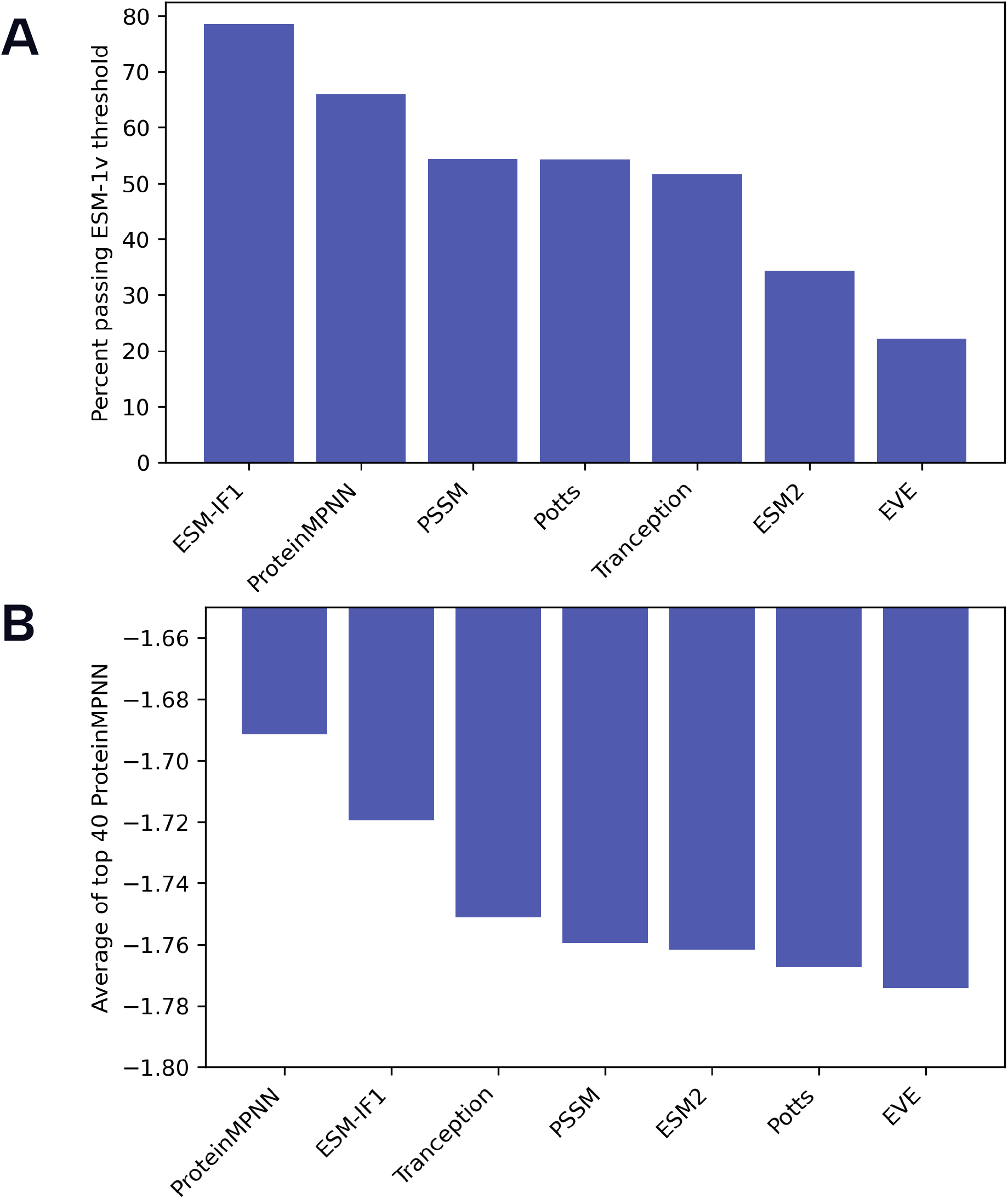
Selection metrics from (Johnson et al., 2024) provide a baseline model ranking. (A) Percentage of generated sequences passing the ESM-1v threshold, defined as exceeding the 90th percentile of scores from natural homologous sequences. (B) Average ProteinMPNN score of the top 40 sequences per model, where sequences are first filtered by the ESM-1v threshold in (A), then ranked by ProteinMPNN score, and the top 40 are averaged. Together, these metrics provide a baseline for ranking models and suggest strong performance of structure-based approaches.

**Figure S3.**
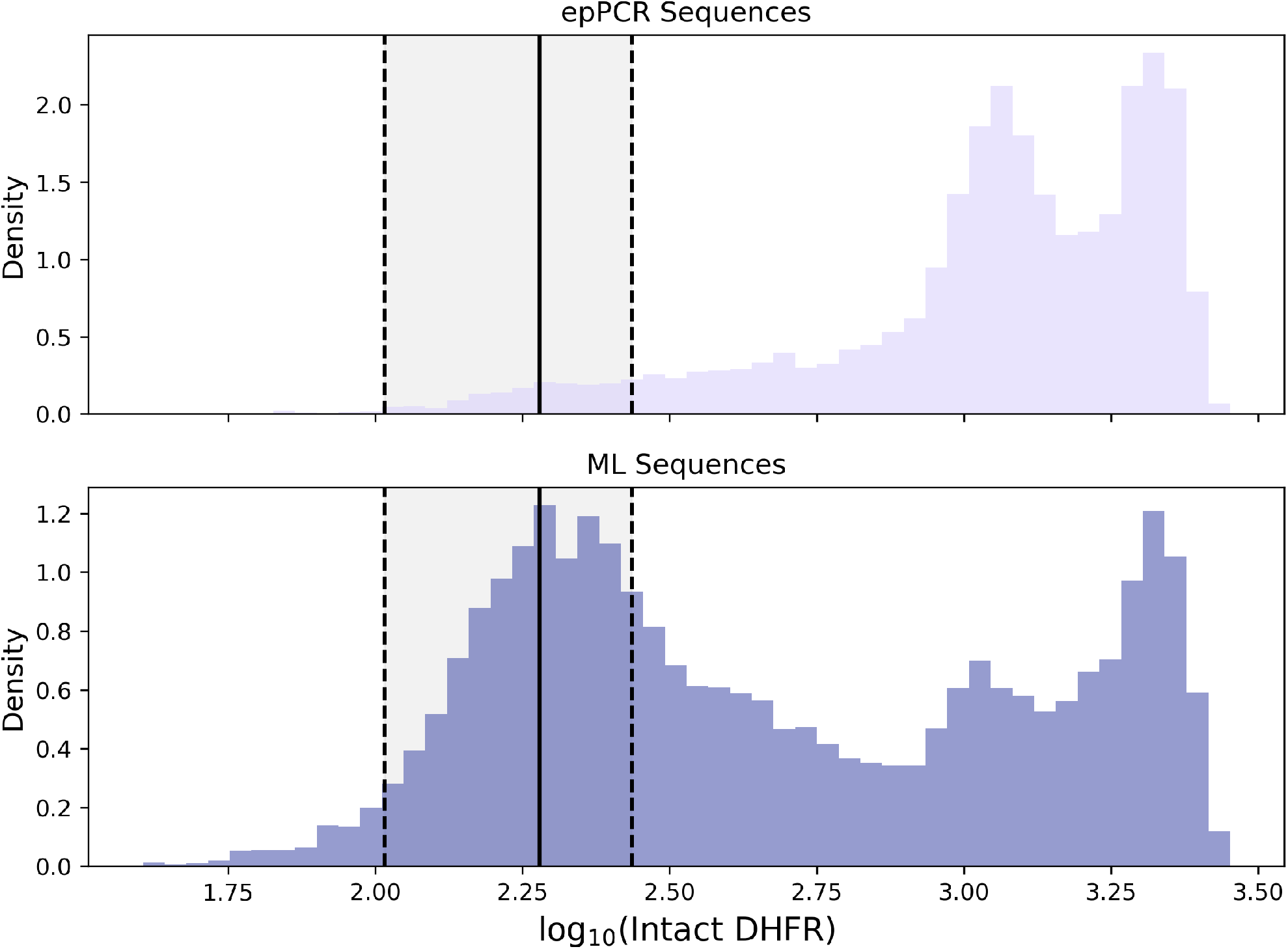
ML-designed libraries enrich for functional variants compared to random mutagenesis. Distributions of experimental measurements (log_10_(Intact DHFR), lower = higher protease activity) for epPCR and ML-designed sequences. The solid line indicates mean wildtype activity, and dashed lines denote the highest and lowest wildtype replicates defining hit thresholds. ML-designed sequences show a higher hit rate (39.32%) than epPCR (6.06%), despite similar mutational loads.

**Figure S4.**
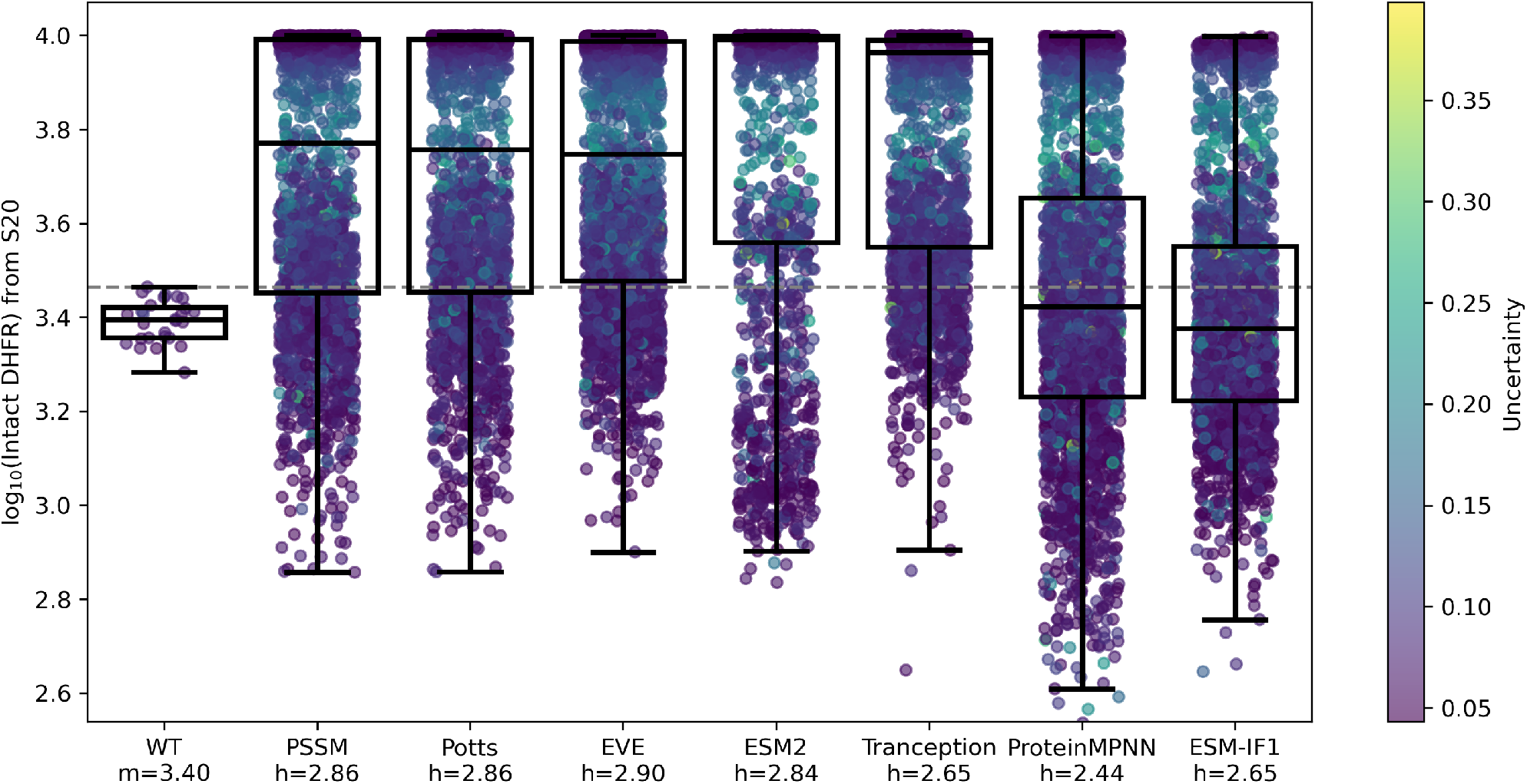
Experimental performance across models under S20 conditions. Distributions of protease activity (log_10_(Intact DHFR), where lower values indicate higher protease activity) measured under the S20 condition, which provides enhanced resolution among top-performing variants. Points are colored by measurement uncertainty. The wildtype distribution is summarized by its median (*m*), while model labels report the best observed variant (*h*) for each model. ProteinMPNN shows substantial enrichment of high-performing variants, with the top variant achieving an ∼9-fold improvement over wildtype. Top variants from Tranception and ESM-IF1 show similar improvements (∼5–6-fold), while ESM2, EVE, PSSM, and Potts exhibit more modest gains (∼3–4-fold). The dashed line indicates the least active wildtype replicate.

## Notes

### Competing Interest Statement

The authors have declared no competing interest.

